# Top-down inputs drive neuronal network rewiring and context-enhanced sensory processing in olfaction

**DOI:** 10.1101/271197

**Authors:** Wayne Adams, James N. Graham, Xuchen Han, Hermann Riecke

## Abstract

Much of the computational power of the mammalian brain arises from its extensive top-down projections. To enable neuron-specific information processing these projections have to be precisely targeted. How such a specific connectivity emerges and what functions it supports is still poorly understood. We addressed these questions in silico in the context of the profound structural plasticity of the olfactory system. At the core of this plasticity are the granule cells of the olfactory bulb, which integrate bottom-up sensory inputs and top-down inputs delivered by vast top-down projections from cortical and other brain areas. We developed a biophysically supported computational model for the rewiring of the top-down projections and the intra-bulbar network via adult neurogenesis. The model captures various previous physiological and behavioral observations and makes specific predictions for the cortico-bulbar network connectivity that is learned by odor exposure and environmental contexts. Specifically, it predicts that after learning the granule-cell receptive fields with respect to sensory and with respect to cortical inputs are highly correlated. This enables cortical cells that respond to a learned odor to enact disynaptic inhibitory control specifically of bulbar principal cells that respond to that odor. Functionally, the model predicts context-enhanced stimulus discrimination in cluttered environments (‘olfactory cocktail parties’) and the ability of the system to adapt to its tasks by rapidly switching between different odor-processing modes. These predictions are experimentally testable. At the same time they provide guidance for future experiments aimed at unraveling the cortico-bulbar connectivity.

**Author summary:** In mammalian sensory processing, extensive top-down feedback from higher brain areas reshapes the feedforward, bottom-up information processing. The structure of the top-down connectivity, the mechanisms leading to its specificity, and the functions it supports are still poorly understood. Using computational modeling, we investigated these issues in the olfactory system. There, the granule cells of the olfactory bulb, which is the first brain area to receive sensory input from the nose, are the key players of extensive structural changes to the network through the addition and also the removal of granule cells as well as through the formation and removal of their connections. This structural plasticity allows the system to learn and to adapt its sensory processing to its odor environment. Crucially, the granule cells combine bottom-up sensory input from the nose with top-down input from higher brain areas, including cortex. Our biophysically supported computational model predicts that, after learning, the granule cells enable cortical neurons that respond to a learned odor to gain inhibitory control of principal neurons of the olfactory bulb, specifically of those that respond to the learned odor. Functionally, this allows top-down input to enhance odor discrimination in cluttered environments and to quickly switch between odor tasks.

## Introduction

A key property of the mammalian brain that is essential for its vast computational power is the pervasiveness of centrifugal, top-down feedback from higher to lower brain areas [1]. The top-down projections can provide the receiving brain area with information that is not available in the feedforward stream [2] and can switch the processing by the lower brain area between different modes [3]. From a theoretical perspective, it has been posited that top-down signals can direct the lower brain area to suppress response to expected inputs or to inputs that have already been recognized by the higher brain area [4–7] and to transmit predominantly information about unexpected or unexplained inputs and the error in the prediction of the inputs [8], focusing on task-relevant information. Such specificity in the processing requires that the top-down projections be precisely targeted [6,9–12]. The mechanisms that are at work in the formation of these specific connectivities are, however, not well understood. Here we address this issue in the context of the extensive structural plasticity of the adult olfactory system.

In the olfactory bulb, which is the first brain area receiving sensory input from the nose, structural plasticity is not restricted to early development, but is also pronounced in adult animals. At that point its key players are the granule cells (GCs), which receive sensory input from the bulb’s principal neurons - mitral and tufted cells (MCs)-and which are the target of massive top-down projections from the olfactory cortex [13–15]. Not only do the GC dendritic spines exhibit strong and persistent fluctuations [16, 17], but these interneurons themselves, which constitute the dominant neuronal population of the bulb, undergo persistent turnover through adult neurogenesis [18]. Both, spine fluctuations and the survival of the granule cells depend on the sensory environment [16, 19]. Moreover, the receptive fields of the adult-born GCs evolve over time [20], reminiscent of the changes in the connectivity between MCs and GCs that occur during learning [21].

Functionally, adult neurogenesis is observed to enable and improve various aspects of learning and memory [22,23]. A particularly clear example is the perceptual learning of a spontaneous odor discrimination task [24]. The survival of GCs seems also to play an important role in retaining the memory of an odor task, with the survival of odor-specific GCs being contingent on the continued relevance of that odor memory [25]. This suggests that GCs receive non-sensory, task-related information via the top-down projections. These projections likely also underlie the specific GC activation patterns that can be evoked by context alone, without the presence of any odor [26].

Taken together, the experiments suggest that sensory input together with non-olfactory information like context or valence may shape the connectivity within the olfactory bulb as well as that of the top-down projections onto the GCs. It is, however, poorly understood how the bottom-up and top-down inputs into the GCs jointly shape the network connectivity [27], what mechanisms are at work, what kind of connectivities arise, and what kind of functionalities these connectivities support. To address these issues we have employed computational modeling using a biophysically supported framework. This model makes a number of predictions that can be tested with physiological and behavioral experiments and that can guide future experiments aimed at unraveling the cortico-bulbar connectivity.

**Fig 1.**
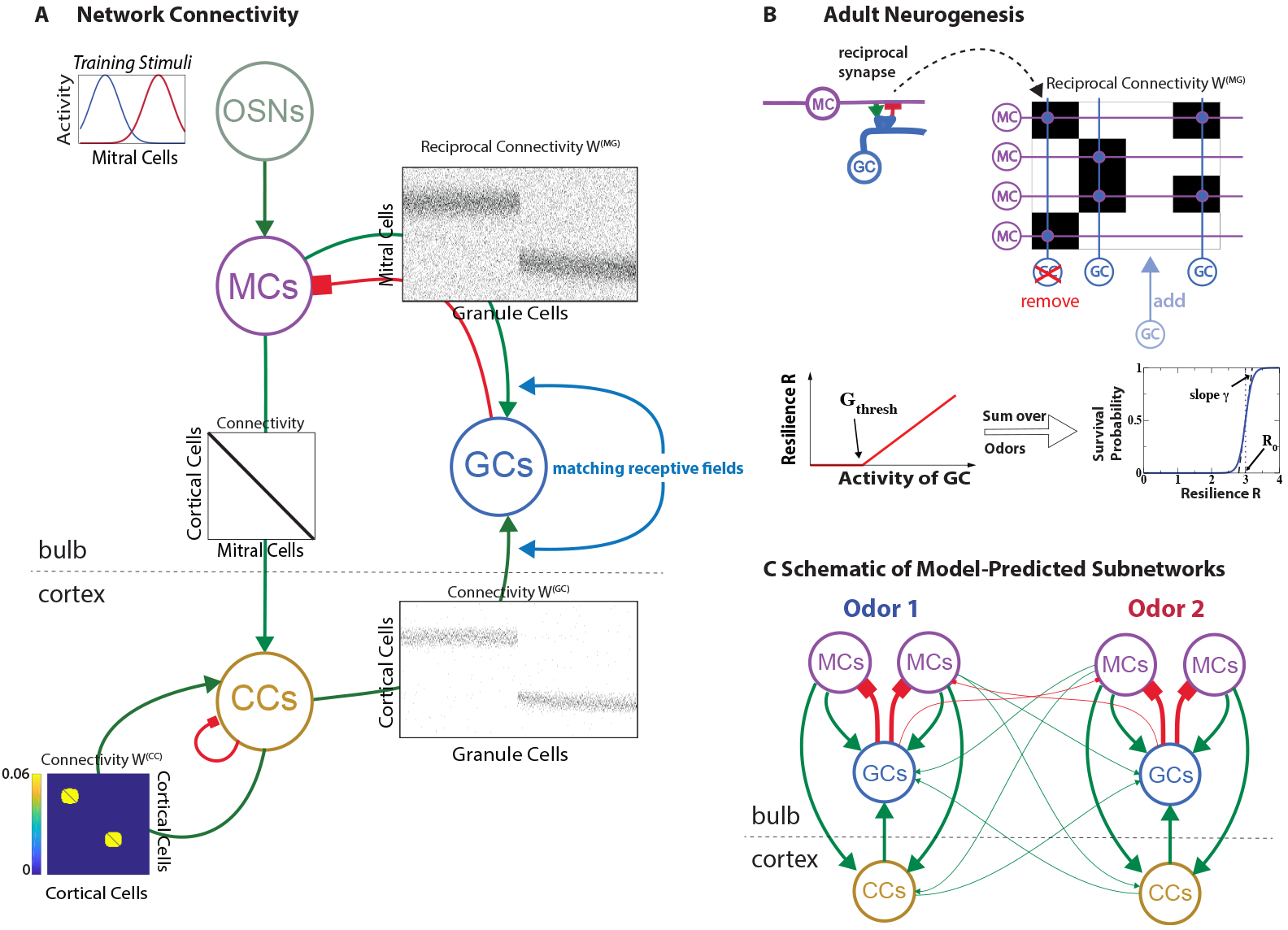
Computational Model. A) Network structure emerging after learning 2 training stimuli. The modeled neuronal populations are indicated by circles marked OSNs (sensory neurons), MCs, GCs, and CCs. Excitatory connections in green with arrows, inhibitory ones in red with squares. Connectivity matrices indicate the connectivities between the individual neurons of the populations (black=connection, white=no connection; cf. Fig.1B). For the resulting disynaptic mutual inhibition of MCs see supplementary Fig.S1 Fig. For the intra-cortical connectivity *W*^(*CC*)^ the colors indicate the synaptic strength of the connections. B) Neurogenesis adds GCs, which make random reciprocal synapses with MCs and receive excitatory projections from random CCs (not shown). The synapses between the 4 individual MCs and GCs are indicated by black rectangles in the connectivity matrix. GC survival depends on GC resilience summed over the training stimuli. C) Idealized schematic of the network structure emerging after learning two odors: CCs disynaptically inhibit predominantly those MCs that respond to the same odor as the CCs. Line thickness indicates the number of connections.

## Results

We have developed a computational network model for the olfactory bulb and its communication with a cortical area, which is based on key features of adult neurogenesis as observed experimentally. Details of the model are given in *Methods*. Briefly, the bulbar component of the model network comprised the two main neuronal populations of the olfactory bulb, the mitral/tufted cells (MCs) and the granule cells (GCs). A third population of principal, excitatory cells (CCs) received inputs from the MCs and projected back to the GCs, thus capturing essential aspects of the top-down projections. The CCs were not meant to model a specific neuronal population in a specific higher brain area; instead, they were intended to represent a neuronal population that mimics certain aspects of the various components of olfactory cortex. For simplicity, we call these principal cells ‘cortical cells’. In Figure 1A each population is indicated by a circle and the connections between the individual neurons of the various populations are shown in terms of connectivity matrices (black=connection, white=no connection between the respective neurons as illustrated in Fig.1B). The MCs received excitatory input from the sensory neurons (OSNs) and formed excitatory projections to the CCs as well as reciprocal synapses with the inhibitory GCs. To aid visualization of the results each CC received input from only one MC, resulting in a diagonal connectivity matrix. The CCs formed excitatory synapses onto the GCs. In addition, they had all-to-all recurrent excitatory connections among themselves, which were endowed with Hebbian plasticity to give this network autoassociative properties [31,32], and inhibited each other via an unmodeled interneuron population.

All neurons were described using nonlinear firing rate models. The MCs and GCs had threshold-linear firing-rate functions. The MCs exhibited some spontaneous firing in the absence of odor input, while the GCs responded only when their combined input from MCs and CCs surpassed a positive threshold. The firing-rate function of the CCs was taken to be sigmoidal.

The key aspect, network restructuring by adult neurogenesis, was implemented by persistently adding and removing GCs. Newly added GCs formed randomly chosen connections with a subset of the MCs and of the CCs. Figure 1B depicts only the reciprocal connections between GCs and MCs, overlaid onto the corresponding connectivity matrix. GCs were removed with a probability that decreased sigmoidally with their ‘resilience’. The resilience of a GC was taken to be the sum of its thresholded activities, which were driven by MCs and CCs in response to a set of training stimuli. This activity-enhanced survival of GCs was motivated by observations in enriched and deprived environments [19,33] as well as experiments with genetically modified GCs [34].

The network structure arising in this model for a range of suitable parameter values is illustrated in Fig.1A for a simple set of two training stimuli that activated partially overlapping sets of MCs. Throughout we used such simplified odor-evoked patterns rather than natural stimuli [17,35], because they facilitated the visualization and understanding of the emerging network connectivities. Since all connections were learned based on activities, the spatial arrangement of the various neurons played no role in the model. Exposing the network to these odors lead to a strengthening of the recurrent connections among the CCs that were co-activated by the odors, implementing an associative memory of these odors. Due to the non-zero threshold for GC activation, for a GC to survive it needed to be co-activated by multiple MCs and CCs at least for some of the training stimuli. Thus, the surviving GCs were connected predominantly to MCs and CCs that had similar receptive fields.

To wit, in Fig.1A one population of GCs was mostly driven by MCs and CCs responding to the left training stimulus, while the other population responded to the one on the right. Due to the reciprocal nature of the MC-GC synapses [36] this induced disynaptic inhibition of MCs not only by other MCs with similar receptive fields [17, 35], but also by cortical CCs that share that receptive field (cf. supplementary Fig.S1 Fig). Thus, the emerging network structure is characterized by subnetworks or network modules, each of which is associated with a learned odor and provides a bidirectional projection between the bulbar and cortical representation of that odor. This is idealized in Figure 1C, where the circles again represent neuronal populations and the thickness of the lines indicates the number of neurons involved in that type of connection. Through this cortico-bulbar feedback structure cortical cells have inhibitory control specifically over those MCs that provide their dominant sensory input. Of course, depending on the similarity of the odors in the training set their associated subnetworks are more or less overlapping or intertwined.

The simulation experiments presented in this paper are for one set of model parameters. Extensive further analysis has shown, however, that our results, particularly for the network structure, are robust with respect to parameter changes.

What ramifications does this odor-specific network structure have for stimulus processing? In the following we use simulation experiments to address this question in a number of behaviorally relevant settings.

### Perceptual Learning of Odor Discrimination

Behavioral experiments on spontaneous odor discrimination using a habituation protocol have shown that exposure to an odor related to the odors used in the discrimination task can induce a perceptual learning of that task; however, this learning was compromised when adult neurogenesis was suppressed [24]. A parsimonious interpretation of this finding is that the restructuring of the bulbar network by adult neurogenesis enhances differences in the bulbar representations of the similar odors rendering them more discriminable. To assess the impact of the network structure on odor discriminability we envisioned a read-out of the bulbar output that consists of the sum of the suitably weighted outputs of all MCs. Discriminability can then be characterized by Fisher’s linear discriminant 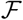 given by the square of the difference between the trial-averaged read-outs corresponding to the two odors divided by the trial-to-trial variability of the read-outs. Our firing-rate framework did not include any trial-to-trial variability. We therefore took as a proxy for it the firing rate, which would be proportional to the variability if the rates arose from Poisson-like spike trains. We considered here the optimal value 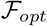 that is obtained if the weights of the outputs to the read-out are chosen to maximize 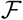 for the stimuli in question. Such optimal weights could be the result of the animal learning the task. For similar odors 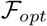 typically increased in our model as the network structure evolved in response to these odors, typically in parallel with a reduction in the Pearson correlation of the MC activity patterns (cf. [35]), capturing the observed perceptive learning [24].

### Odor-specific Apoptosis after Extinction of Odor Memory

The survival of the model GCs depended on the sensory input they received from the MCs as well as on the top-down input from the CCs. The top-down input was enhanced if the presented odor had previously been ‘memorized’, i.e. if the corresponding recurrent excitatory connections among the CCs had been strengthened. What happens if the cortical memory of one of the learned odors is erased, i.e. if the recurrent connections of the CCs representing that odor are removed? In our simulations removal of the memory of odor pair 1 (Fig.2B) significantly enhanced cell death (Fig.2C), affecting mostly the GCs that previously had responded to the odors in that pair and whose top-down inputs had been enhanced by its cortical odor memory (Fig.2D and supplementary Fig.S2 Fig). In parallel, the network’s ability to discriminate between the odors in that pair was substantially reduced (Fig.2E). This degradation of the performance did not occur if the removal of GCs was blocked in the simulation.

These results capture essential features of experiments in mice in which the extinction of an odor memory enhanced the apoptosis of GCs, particularly of those GCs that had been responsive to that odor. However, fewer of the GCs died and the mice did not forget the task when apoptosis was blocked during the extinction of the odor memory [25].

**Fig 2.**
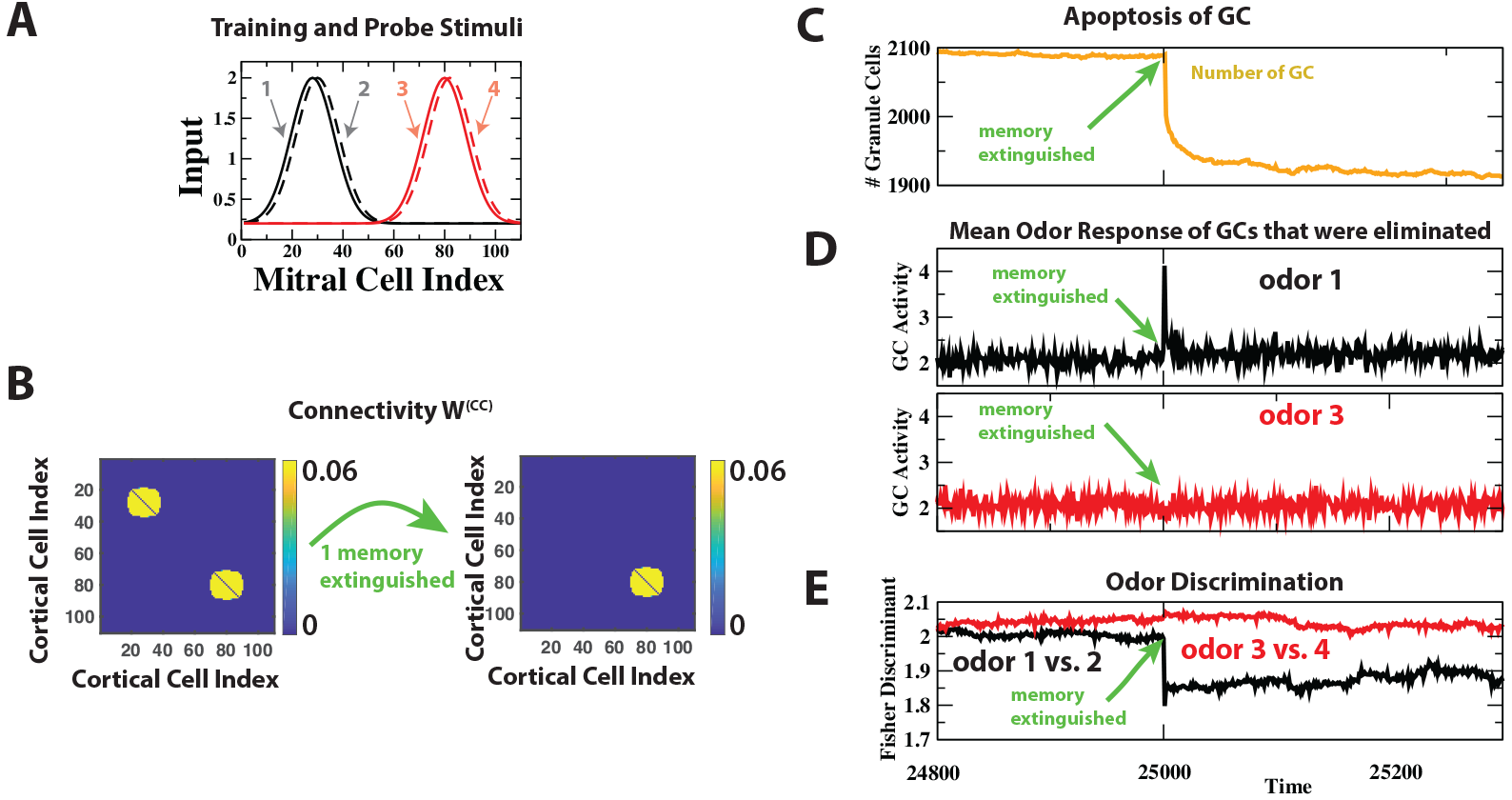
Extinction of cortical odor memory eliminates GCs. A: Training and probe odors. B: Extinguishing the cortical memory of odors 1 and 2 eliminated the corresponding cortical associational connections. C: Extinction induced apoptosis of many GCs. D: The GCs that were removed in the extinction had strongly responded to odor 1 before the extinction, but not to odor 3. E: The extinction-induced reduction in inhibition compromised the discrimination between odors 1 and 2, but not between odors 3 and 4.

### Specific GC Activation in the Absence of Odor Stimulation

Experimentally it has been found that specific GC activity patterns could be evoked even in the absence of odors, if the animal was placed in an environment that previously had been associated with an odor [26]. In fact, the GC activation pattern that was induced by this environment had substantial overlap with the pattern evoked by the associated odor. This was not the case for a different environment. A natural interpretation of these observations is that the odorless GC activation patterns were driven by top-down projections onto the GCs from higher brain areas that have access to non-olfactory, e.g. visual, information [26]. To capture this aspect we extended our cortical model to include CCs that did not receive direct input from the olfactory bulb but were driven by non-olfactory, contextual information. This could, for instance, represent information from other sensory modalities, information about the task the animal is to perform, or an expectation by the animal. We introduced excitatory associational connections with Hebbian plasticity between these cells and the CCs that received MC input and extended the global inhibition to those cells.

Presenting two odors to the network, each in the presence of a different context, established associational connections in the cortical network between the CCs that received odor input (cell indices 1 to 110) and the CCs that received the corresponding contextual input (marked’context 1’ and’context 2’ in Fig.3B). This enabled the contextual input to excite GCs via top-down projections (supplementary Fig.S1 FigB) even in the absence of any odor stimulation. Due to the specificity of the cortico-bulbar network the resulting odorless GC activity patterns were highly correlated with the patterns induced by the associated odor, but not with those of the other odor (Fig.3C,D), recapitulating the experimental observation. In contrast, the context-evoked, odorless MC patterns were anti-correlated with the odor-evoked patterns, akin to representing a novel ‘negative’ odor, which may evoke a very different percept than the odor itself. This may account for the enhanced response of the animals observed in the familiar, but odorless context, but not in the unfamiliar context [26].

**Fig 3.**
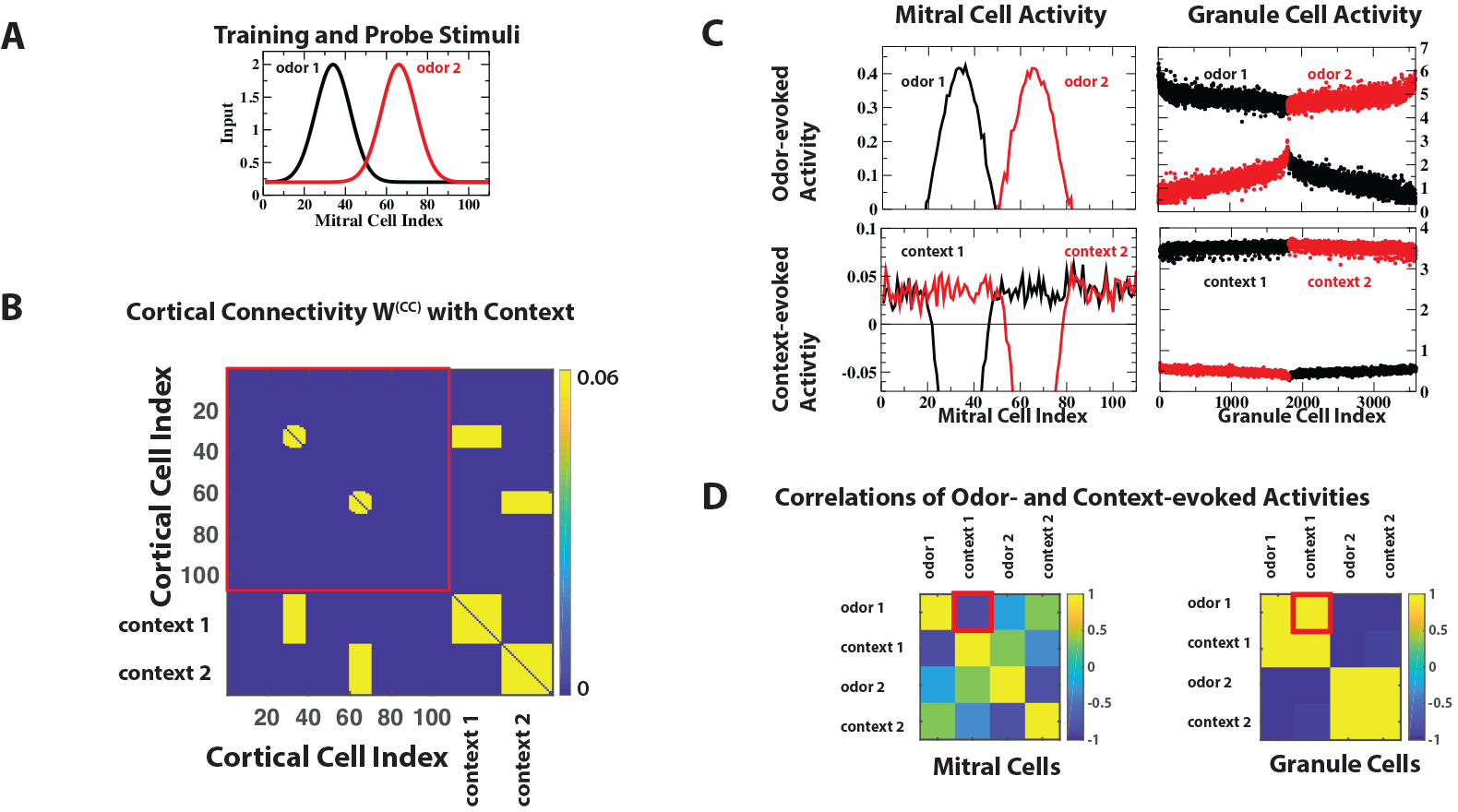
Context induces specific GC activity. A,B: The 2 training stimuli were associated with contexts 1 and 2, respectively. C,D: Context-and odor-evoked MC activities are anti-correlated, while context-and odor-evoked GC activities are highly correlated (red boxes in D).

What functionality is enabled by the learned network structure, which allows CCs to inhibit specific MC? To assess this question we considered two scenarios: i) the detection and discrimination of odors in a cluttered environment and ii) rapid switching between different odor tasks.

### Context Enhances Detection and Discrimination in Cluttered Odor Environments

We considered cluttered environments in which the presence of additional odors may or may not occlude (mask) the odors of interest, an olfactory analog of the ‘cocktail party’ problem [37]. As an illustrative example we considered the detection of a weak target odor. In the presence of a strong odor that activated a large number of MCs and occluded the target odor this required the discrimination between the occluder alone and the occluder with the target (Fig. 4E). This was difficult because the MCs carrying the information about the target were also driven by the occluder, rendering the relative difference between the overall pattern with target (red, solid symbols) and that without target (black, open symbols) small. If the activation by an occluding odor could be reduced without suppressing the contribution from the target odor, detection of the target should be significantly enhanced. Indeed, in our model such an odor-specific inhibition was possible if the occluding odor was familiar, i.e. if it had been one of the training stimuli.

In these simulation experiments we associated the occluding odor during the training with context 1 (Fig.4D). The odor-specific inhibition had then two contributions. The intra-bulbar component did not require cortical activation and suppressed the familiar occluder on its own (Fig.4F left panel). In the presence of the context the detectability of the target was even further enhanced by the cortical excitation of the GCs (Fig.4G left panel), increasing the Fisher discriminant 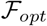(Fig.4H). However, the same context was detrimental for the detection of the target odor if the occluding odor was not present: the strong context-enhanced inhibition almost eliminated the response to the odor (Fig.4F,G right panels). The flexibility in the control afforded by top-down inputs therefore substantially enhanced performance.

**Fig 4.**
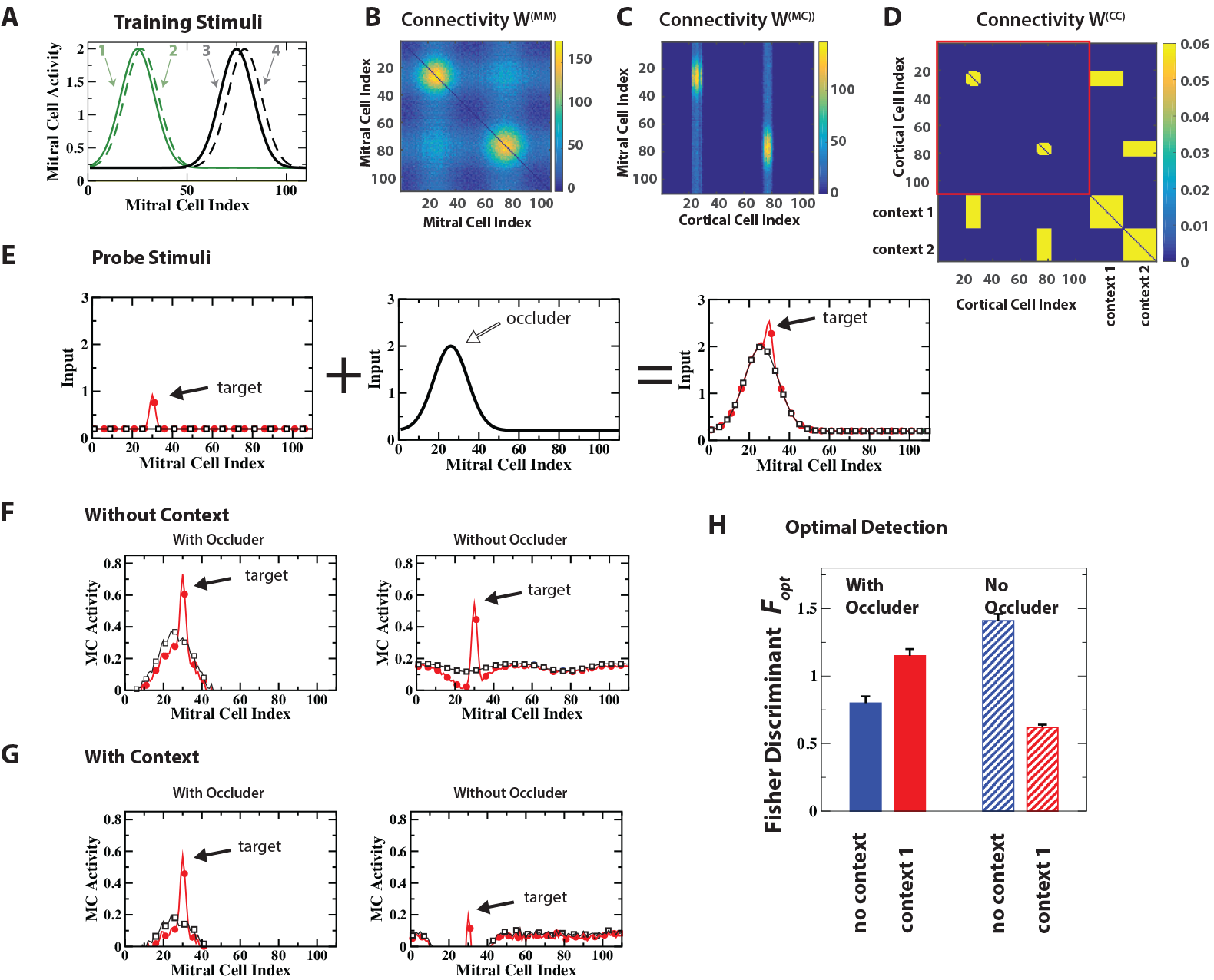
Context-enhanced Odor Processing in Cluttered Environments: Occluding stimulus. A,D: Training stimuli 1,2 and 3,4 were associated with contexts 1 and 2, respectively. B,C: learned disynaptic inhibitory connections among MCs and from CCs to MCs. E: Probe stimuli consisted of a weak target odor with or without stimulus 1 as a strong occluding odor. F,G: MC activities resulting from probe stimuli with and without context 1. H: Context 1 increased the Fisher discriminant 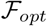 when the occluding odor was present, but reduced 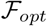 when it was absent.

If the odors of interest are not occluded by any other odors in the environment, read-out cells can, in principle, adapt the weights of their input synapses so as to focus only on the relevant odors and ‘ignore’ the cluttered environment. However, this weight optimization is often not possible, since the animal may not know yet which odors are to be discriminated. This is, for instance, likely the case in the early phase of learning a new discrimination task [38]. During this phase it is reasonable to envision that animals rely on a large number of read-out cells each of which receives inputs from different combinations of MCs and is therefore sensitive to different aspects of the MC activity patterns. The activity of many of these read-out cells will be dominated by the uninformative components of the odor environment (‘distractor’), making it difficult to discriminate between two weak target odors, even if they are not occluded by the environment (Fig.5E, right panel illustrating optimal and random read-out). To analyze such a situation we employed a non-optimal Fisher discriminant 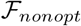 based on a large number of random read-outs of the MCs (cf. [10] in the Methods).

**Fig 5.**
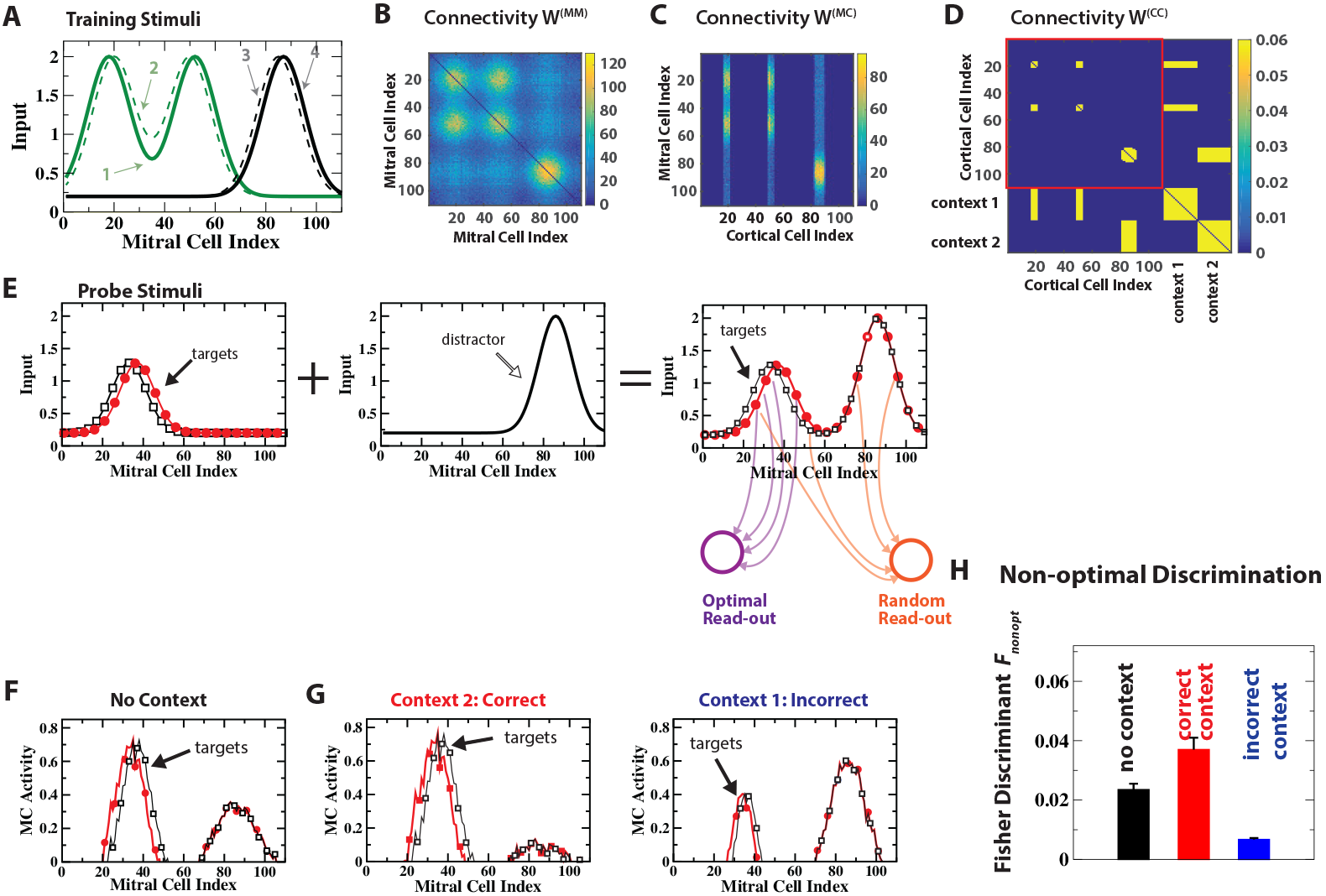
Context-enhanced Odor Processing in Cluttered Environments: Distracting stimulus. A,D: Training stimuli 1,2 and 3,4 were associated with contexts 1 and 2, respectively. B,C: learned disynaptic inhibitory connections among MCs and from CCs to MCs. E: the probe stimuli consisted of one of the weaker target odors combined with training stimulus 3 as a strong distractor. Sketch of connections for an optimal read-out that focuses on the target odors and for a non-optimal, random read-out. F,G: MC activities with and without context. H: The correct context suppressed the distractor and enhanced the Fisher discriminant 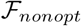 of the random read-out. In contrast, the incorrect context reduced 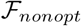.

We considered the discrimination between two similar, novel target odors in the presence of a strong, familiar odor that did not occlude the targets but served as a distractor (Figure 5E). It was associated with context 2. In addition, the network was familiarized with odors 1 and 2, which were associated with context 1 and partially overlapped with the novel target odors (Figure 5A-D). Even in the absence of any contextual signal the learned intra-bulbar connectivity was able to suppress to some extent the distracting, familiar odor relative to the novel odors. In the presence of context 2, which was associated with the distractor (’correct’ context), this suppression was substantially enhanced through the cortical feedback driven by that context, leading to much better discriminability of the two novel odors (Figure 5G,H).

As in the case of discrimination via an optimal read-out (Fig.4) it was not beneficial to have indiscriminate strong cortical feedback for all familiar odors, even if these odors were not present. For instance, the feedback driven by context 1 (’incorrect’ context) was detrimental to the discrimination of the target odors if neither of the familiar odors 1 and 2, which were associated with context 1, were part of the odor scene. While the target odors were not occluded by these familiar odors, they had significant overlap with them. Therefore the connectivity that was learned through the training included a sizable number of inhibitory projections - via the GCs - from the CCs representing context 1 to the MCs representing the target odors. This lead to a strong suppression of the MC response to the target odors, resulting in poor discriminability (Figure 5G,H).

Thus, the ability of top-down inputs to induce specific inhibition in a *flexible* manner substantially enhanced olfactory processing.

### Top-Down Input Enables Task Switching

The neurogenic evolution of the structure of the cortico-bulbar network occurs on a time scale of days, particularly since the apical reciprocal MC-GC synapses form only days after the proximal synapses of the top-down projections [39]. Synaptic plasticity based on changes in the synaptic weights can act on much shorter time scales, allowing cortical associational connections to adapt faster to changes in the tasks that the animal needs to perform. This could modify the top-down signals to the bulb, altering bulbar processing.

**Fig 6.**
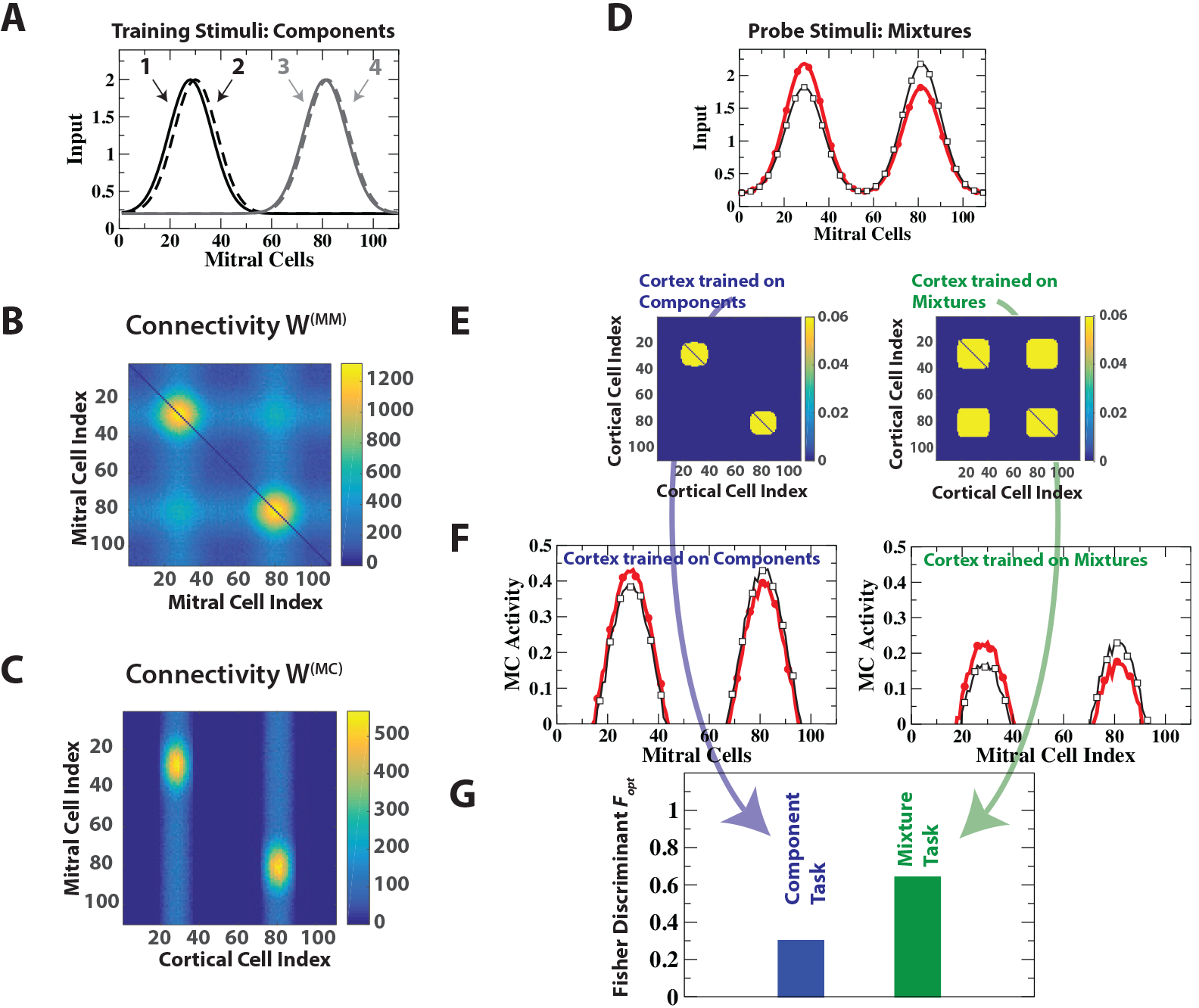
Cortical task switching enhances discrimination. A: training stimuli consisted of 2 pairs of dissimilar stimuli. B,C: resulting bulbar and cortico-bulbar connectivity. D: probe stimuli consisted of two mixtures of the training stimuli. E: cortical connectivities after cortical training on the individual components (left) or the mixtures (right). F,G: MC activities and optimal Fisher discriminant 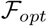 with cortex trained on the components or the mixture, respectively.

As an example we considered a situation in which the network was trained on two pairs of odors, *O*_1,2_ and *O*_3,4_. The two odors in each pair were very similar, but the pairs were dissimilar from each other (Figure 6A). As expected, the training increased the discriminability of the odors within a pair. However, for the discrimination of the mixture *M*_1_ = 0.55 *O*_1_ + 0.45 *O*_3_ from the mixture *M*_2_ = 0.45 *O*_1_ +0.55 *O*_3_, obtained by combining odors from the two pairs, the training on the individual components *O*_1,2_ and *O*_3,4_ was detrimental. While it established mutual inhibition of MCs that were activated by the *same* mixture component (*O*_1_ *or O*_3_), it provided only little mutual inhibition between MCs that were activated by different mixture components: the number of connections between MCs representing odor *O*_1_ (MCs with index near 30 in Fig.6B) and MCs representing odor *O*_3_ (MC index near 80) was small. However, these MCs are activated simultaneously in the mixtures and mutual inhibition of these MCs is needed to enhance the discrimination between the mixtures [17,35,40,41]. As a result the inhibition significantly reduced the relative difference between the two mixtures and with it their Fisher discriminant (Figures 6F left,G). Inhibition between the MCs representing *O*_1_ and those representing *O*_3_ can be effected by establishing associational excitatory connections between the CCs that disynaptically inhibit these two groups. To do so we exploited the Hebbian plasticity of the cortical synapses and trained the cortical network briefly to the mixture 0.5 *O*_1_ +0.5 *O*_3_ (Figure 6E right). This enhanced the discriminability of the mixtures substantially (Figure 6F right,G).

Thus, cortical projections can exploit the learned, odor-specific subnetwork structure (Fig.1C) to switch between different cortico-bulbar processing modes, adapting to the odor objects at hand.

### Experimental Predictions

The key anatomical feature of the model network resulting from learning is its connectivity. Specifically, the projections that the GCs receive from the MCs and from the CCs are predicted to be matched: a given GC receives cortical inputs predominantly from those CCs that respond to the same odors as the MCs projecting to that GC. This matching of the GCs’ receptive fields can be tested experimentally. One possibility is to express-after suitable training-ChR2 conditionally (e.g., via c-Fos [42]) in those principal cells of piriform cortex that are activated by the training odor, combined with expression of a calcium-indicator in the GCs. Note that the training needs to activate the neurogenic plasticity of the bulb [24], which is not the case if the odor exposure is only passive [43]. The model predicts that optical stimulation of cortical cells in the absence of an odor will then lead to excitation patterns of the GCs that are strongly correlated with the patterns excited by the training odor (Fig.3C,D). Previous experiments in which odor-evoked and context-evoked GC activation patterns were found to be correlated in different animals are suggestive of this outcome [26].

On a behavioral level the model makes specific predictions for the learning of odor discrimination or detection in cluttered environments. Recent experiments have shown that the detection of an odor in a go/no-go task is particularly difficult if that odor is masked or occluded by an odor [37]. Our model predicts that the performance in such a detection task would be enhanced if the animal is first familiarized with the occluding odor over an extended period of time [24]. The resulting restructuring of the bulbar network would lead to a reduction in the response to that familiar odor, partially unmasking the task-relevant odor. If, in addition, the occluding familiar odor has been associated with a non-olfactory context [26], the model predicts that the performance is further enhanced if the task is performed in that context, but not in a different, novel context (Figure 4A).

Even if the cluttered odor environment does not occlude the task-relevant odors, it is expected that the learning speed in an odor-discrimination task [38] is reduced by strong distracting odors. The model predicts that sufficient familiarization [24] with the distracting odors will reduce their uninformative contributions to the overall MC activation pattern. This is expected to increase the signal-to-noise ratio and with it the learning speed by reducing the contributions from the uninformative MCs to the variability of the read-out. If, in addition, the distracting, uninformative odors are associated with a context, the learning speed is predicted to increase further in the presence of that context (Figure 5B). If the cluttered environment makes the task too hard to learn for naive animals, our model suggests that prior familiarization with the distracting odors, preferably in a specific context, may enable the animals to master this difficult task.

## Discussion

Adult neurogenesis is a striking mechanism of structural plasticity that has the potential to rewire a network extensively. In mammals it arises predominantly in two brain areas. In the dentate gyrus it involves excitatory granule cells; their role in the network has been studied in detail [44,45], also in terms of computational models [46]. In the olfactory system, where adult neurogenesis involves inhibitory rather than excitatory granule cells, there is also a substantial body of experimental work [18], but only few modeling studies are available [35, 47]. Here we have developed a computational model for the neurogenic evolution of the network connectivity with an emphasis on the possible role of the pervasive top-down projections from cortical areas. The model is based on a number of experimental observations: GC survival depends on GC activity [19,34], GC activity can be induced in the absence of odor stimulation [26], and piriform cortex exhibits extensive recurrent excitation, which can support associational memory [31,32]. The model captures qualitatively the experimentally observed perceptual learning afforded by neurogenesis [24] as well as the enhanced apoptosis of specific GCs and the reduced odor discriminability after the extinction of memories [25].

Without theoretical guidance, it is difficult in experiments to identify the functional structure of the cortico-bulbar connectivity, in particular, because odor representations in the olfactory system do not reflect detailed spatial maps like those of other sensory systems [48, 49]. An important contribution of the model is therefore its prediction that through the structural plasticity the network develops a structure that reflects the learned odors and provides enhanced inhibition that is specific to these odors. This inhibition is in part intra-bulbar and in part driven by top-down (cortical) inputs. The latter reflects the formation of a bidirectional connection between the bulbar and the cortical representation of the familiar odors, which allows the cortical cells associated with such an odor to inhibit specifically those MCs that are excited by that odor. This inhibition is mediated by GCs. For this connectivity to arise the reciprocal nature of the MC-GC synapses is essential. Our results therefore suggest that a key function of the reciprocity of these synapses may be to guide the wiring of the cortico-bulbar network connectivity. The predicted matching of the GC receptive fields for olfactory and for cortical input can be tested experimentally. Moreover, the model can guide future experiments aimed at elucidating the cortico-bulbar connectivity.

Functionally, the model predicts, in particular, that the learned connectivity improves the detection and discrimination of novel odors in cluttered environments, if the occluding or distracting odors are familiar. Moreover, this improvement can be enhanced by top-down input when the occluding or distracting odor activates cortical memory, which may indicate that a higher brain area has recognized parts of the odor scene. This may allow something akin to the ‘explaining away’ of components of a complex odor mixture that is theoretically predicted for the optimal processing of stimuli [4–6,12]. Recent work has identified networks that demix familiar odors employing approximate optimal Bayesian inference; the anatomical structure of these networks is very close to that emerging naturally in our neurogenic model [6, 7]. By providing a biophysically supported mechanism through which the system can learn the required network structure our model complements this abstract normative approach.

Going beyond the purely olfactory aspect, the top-down input could encode task-related expectations or contextual information originating from other sensory modalities. Thus, it may implement a predictive coding in the bulb that reflects the context or task at hand [50]. Our model demonstrates how such contextual information can enhance performance.

The formation of the bulbar and cortico-bulbar network via structural plasticity is a relatively slow process. However, our model predicts that the network structure emerging from it can be exploited by faster synaptic learning processes in cortex, which allow the system to switch relatively quickly between different discrimination tasks. This is reminiscent of the task-dependent switching of neuronal responses observed in V1 [3].

Focusing on the slow evolution of the network structure our model is intentionally minimal with respect to the dynamics of the individual neurons. We describe the neurons in terms of their firing rate and focus on neuronal steady-state activities. So far, the knowledge about the biophysical mechanisms controlling GC survival and their dependence on neuronal activity is not sufficient to guide the development of a detailed model of that process, which would, e.g., connect GC spiking or calcium-levels with GC survival [34,51,52].

To assess the discriminability of MC activity patterns we assumed that the firing rates result from an irregular firing in which the variance of the spike number is proportional, albeit not necessarily equal, to the mean spike number. Thus, we have not taken the widely observed rhythmic aspect of the bulbar activity into account [53], which suggests that animals may also be able to make use of spike-timing and synchrony information in odor processing [5,54,55]. The modular network structure and the associated top-down inputs that are predicted by our model are likely to have also substantial impact on spike timing and synchrony. A study of the resulting dynamics and functional consequences is beyond the scope of this paper and will be left to future work.

The system most likely gains additional richness through the nonlinear response properties of the GCs, which include local dendritic depolarization that can provide reciprocal inhibition to the MCs even without spiking [56], as well as wide-spread dendritic activation associated with calcium-or sodium-spikes that may drive wide-spread lateral inhibition [57, 58]. Thus, it is likely that inhibition can operate in different regimes [59], which may further enhance the system’s ability to switch between different tasks.

Adult neurogenesis is not the only plasticity mechanism operating in the olfactory bulb. Spike-timing dependent plasticity is known to be enhanced in the proximal synapses of young adult-born GCs [14,60]. Long-term depression has been demonstrated at the reciprocal synapses between MCs and GCs [61,62]. In addition, these synapses exhibit substantial structural plasticity in the form of spine fluctuations in adult-born as well as neonatally born GCs [16,17]. In computational models of the olfactory bulb without top-down inputs spine fluctuations and adult neurogenesis lead to very similar network connectivities [17,35]. We therefore expect that a cortico-bulbar model based on spine fluctuations would lead to results that are very similar to the ones based on adult neurogenesis described here. How the two types of structural plasticity complement each other and how the sequential formation of proximal and apical synapses [39] affects the emerging connectivity are interesting questions, which are, however, beyond the scope of this paper.

## Methods

Our model consisted of three populations of neurons, mitral/tufted cells (*M*), granule cells (*G*), and ‘cortical’ cells (*C*). Although the differences between mitral and tufted cells in terms of their properties and function are becoming increasingly known [28,29], the model did not distinguish between them. The neuronal activities were described by the nonlinear rate equations

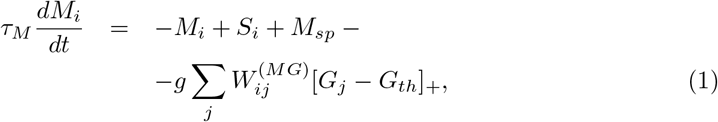

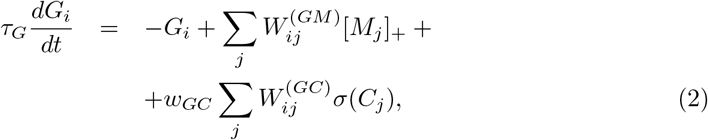

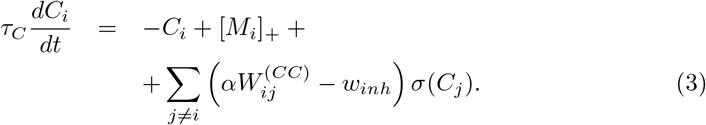

The odor stimuli were given by *S*_*i*_ and the spontaneous MC activity by *M*_*sp*_. The nonlinearity for the MCs and GCs was given by the rectifier

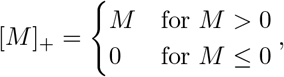

while the nonlinearity for the CCs was sigmoidal

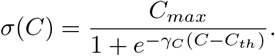

The bulbar connectivity matrices satisfied

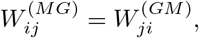

reflecting the reciprocity of the MC-GC synapses. The entries of 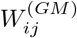 were given by 0 or 1. Without loss of generality the strength of the excitatory synapses from the MCs to the GCs was set to 1, while the strength of the reciprocal inhibition was given by *ɡ*.

The excitatory cortical synapses were taken to be plastic. In all the simulations presented in this paper the cortical connectivity was learned at the beginning of the simulations using the Hebbian plasticity rule

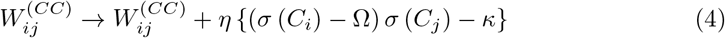

with hard limits given by 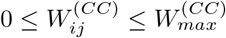 Here, the cortical activities *C*_i_ were given by the steady state reached for the current values of 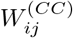. The learning rate was given by *η*, the threshold for potentiation by Ω. There was an overall slow decay of the weights given by *κ*. The parameter *α* in equation (3) allowed a reduction of the recurrent excitation during learning to reduce interference with previously learned memories [30] by switching between *α*_*learn*_ and *α*_*recall*_. During the neurogenic network evolution the strengths of the recurrent cortical connections were held fixed. There was also all-to-all inhibition among the cortical cells with strength *w*_*inh*_.

The connectivities 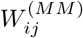 and 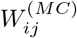 of the effective disynaptic inhibition between MCs and of MCs by CCs, which are shown in supplementary Fig.S1 Fig and Fig.S3 FigB, are given by

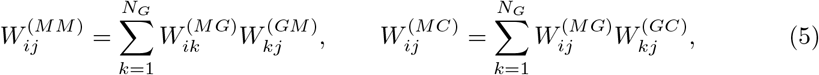

where *N*_*G*_ is the number of GCs. Note that the diagonal elements of 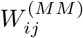 are given by the number of non-zero elements in the corresponding row of 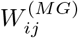 and are typically quite large. In the connectivity figures we have therefore replaced these large values by 0 to reveal the structure of the remaining connections. In the computations these diagonal elements were, of course, not set to 0.

In each time step of the neurogenic network evolution 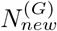 new GCs were added to the network, each making 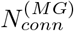 randomly chosen connections with MCs and 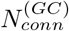 randomly chosen connections with CCs. Then the steady-state values of *M*, *G*, and *C* for each odor 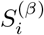, *β* = 1… *N*_*s*_, in the odor environment were determined by solving the evolution equations (1,2,3) for the corresponding odor until a steady-state was reached. Since only the steady-state values were desired we set *τ*_*G*_ = 0, which allowed a drastic reduction of the number of differential equations that had to be solved. Then the resilience of each GC was determined as the sum of its thresholded activity over all *N*_*s*_ training stimuli,

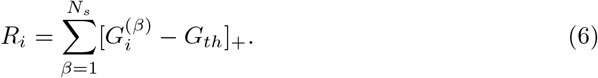

Then GCs were removed depending on their survival probability *P*_*i*_ given by (cf. Figure 1B)

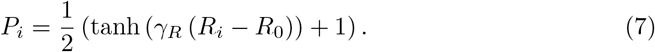

This completed the neurogenic time step. For all results, except in Fig.2, sufficiently many such steps were taken to reach a statistically steady state in the network connectivity and in the various quantities assessing the features of the system.

The number of GCs was not fixed in this neurogenic model; instead, it depended on the strength of the inputs 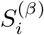 and the strength *ɡ* of the inhibitory connections. For strong inhibition *ɡ* the MC activities were low, leading to low activities of the GCs and a correspondingly low survival probability. As a result, the number of GCs was low. Conversely, choosing weak inhibition *ɡ* lead to a large number of GCs. When the number of GCs was low, adding and removing a single GC had a significant effect on the MC and GC activities, resulting in strong fluctuations in the output of the network. To balance computational effort with the need to keep the fluctuations sufficiently low, we therefore adjusted in the simulations the inhibitory strength *ɡ* during the network evolution to keep the number of GCs within a predetermined target range centered at 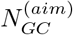. Only in Fig.2, where the focus was the temporal evolution of the network, we kept *ɡ* fixed after *t* = 18, 750 before the memory was extinguished at *t* = 25,000.

Each model odor stimulus was given by *N*_*h*_ combinations of Gaussian activity profiles of the form

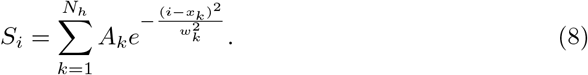

Contextual inputs drove the corresponding contextual cortical cells *C*_*i*_ with a cell-independent amplitude *A*_*context*_.

To assess the discriminability of two stimuli 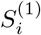 and 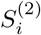 we used the optimal Fisher discriminant *F*_*opt*_

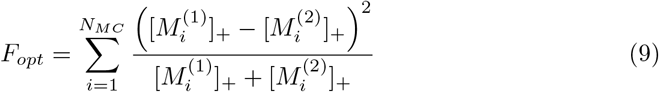

and a non-optimal Fisher discriminant *F*_*nonopt*_, which was obtained as the mean of 5,000 Fisher discriminants *F*_*random*_,

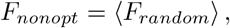

each of which was based on a random read-out of the MC activities,

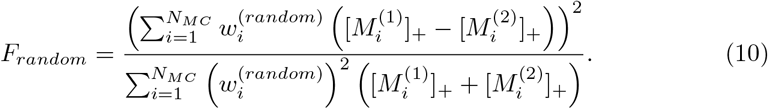

Here the weights 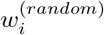 were real numbers between −1 and +1 drawn from a uniform distribution. By including negative weights we allowed for the possibility that the hypothetical read-out cell could also receive disynaptic inhibition from the MCs.

In the simulation experiments presented in this paper we used one set of parameters. We have, however, tested that our results are robust under parameter changes. The parameters used here are as follows:

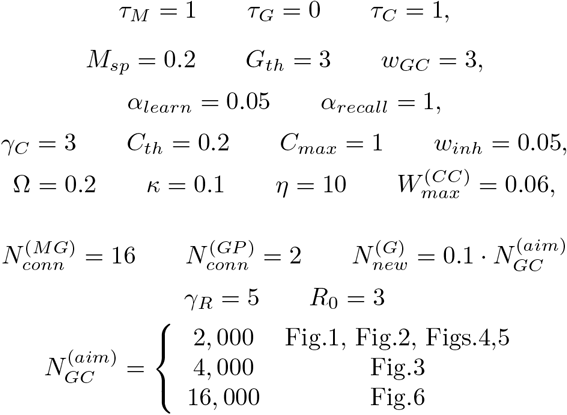

The parameter values for the training stimuli stimuli were

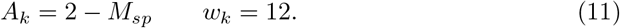

The same parameters were used for the probe stimuli in Fig.2 and Fig.6. In Fig.4 the parameters of the target in the probe stimulus were

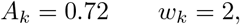

while in Fig.5 the parameters for the targets were

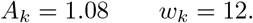

The parameters for the occluder and the distractor in Figs.4,5 were also given by Eq.(11).

The Matlab codes used in these computations are available from the authors upon request.

## Acknowledgments

This research was funded by NIH grant DC015137 (to H.R.). W.A. and J.N.G. received undergraduate support through NSF RTG DMS-0636574. Northwestern University provided HPC access (Quest).

## Supporting information

### S1 Fig. Effective disynaptic inhibition

A) After learning, the GCs effect disynaptic mutual inhibition of MCs that are co-activated by the training odors.

B) The training results in disynaptic inhibition of MCs by CCs that are activated by the same training odors as the MCs (cf. Fig.1A). In both figures the colorbar indicates the number of GCs that mediate the connection between the respective cells.

### S2 Fig. Extinction of a memory

Extinguishing the memory of odor pair 1 removed specifically the GCs that had been associated with that odor pair. A: Connectivity between MCs and GCs before (top) and after (bottom) extinguishing the memory. The number of GCs activated by MCs with index ~30 was reduced by the extinction (cf. arrows). B: Connectivity between CCs and GCs before (top) and after (bottom) extinguishing the memory. The number of GCs excited by CCs with index ~30 was reduced by the extinction (cf. arrows).

### S3 Fig. Top-down connections mediate activation by context

Cortical cells with indices ~30 were activated by context 1 via associational connections (cf. Fig.3B). Through the cortical projections they activated specifically GCs with index below 1805 (A bottom), which in turn inhibited predominantly MCs with index ~30 via the reciprocal connections between MCs and GCs (A top). This resulted in an effective disynaptic inhibition of MCs by CCs (B bottom). Analogously for context 2. Thus, the context excited approximately the same set of GCs as the odor associated with that context. In contrast, the context effectively inhibited the MCs that would be excited by that odor. Due to the reciprocal nature of the synapses the GCs also mediated mutual inhibition between the MCs (B top).

## References

1. Sherman SM, Koch C. The Control of Retinogeniculate Transmission In the Mammalian Lateral Geniculate-nucleus. Exp Brain Research. 1986;63(1):1–20.

2. Restrepo D, Doucette W, Whitesell JD, Mctavish TS, Salcedo E. From the Top Down: Flexible Reading of a Fragmented Odor Map. Trends Neurosci. 2009;32(10):525–531.

3. Li W, Piëch V, Gilbert CD. Learning to link visual contours. Neuron. 2008;57(3):442–451. doi:10.1016/j.neuron.2007.12.011.

4. Ambros-Ingerson J, Granger R, Lynch G. Simulation of Paleocortex Performs Hierarchical-Clustering. Science. 1990;247(4948):1344–1348.

5. Kepple D, Giaffar H, Rinberg D, Koulakov A. Deconstructing Odorant Identity via Primacy in Dual Networks. arXiv:160902202. 2016;.

6. Grabska-Barwinska A, Barthelmé S, Beck J, Mainen ZF, Pouget A, Latham PE. A probabilistic approach to demixing odors. Nature Neuroscience. 2016;20:98. doi:10.1038/nn.4444.

7. Yamada Y, Bhaukaurally K, Madarász TJ, Pouget A, Rodriguez I, Carleton A. Context-and Output Layer-Dependent Long-Term Ensemble Plasticity in a Sensory Circuit. Neuron. 2017;93:1198–1212.e5. doi:10.1016/j.neuron.2017.02.006.

8. Rao RP, Ballard DH. Predictive coding in the visual cortex: a functional interpretation of some extra-classical receptive-field effects. Nat Neurosci. 1999;2(1):79–87. doi:10.1038/4580.

9. Murphy PC, Duckett SG, Sillito AM. Feedback Connections To the Lateral Geniculate Nucleus and Cortical Response Properties. Science. 1999;286(5444):1552–1554.

10. Matsutani S, Yamamoto N. Centrifugal Innervation of the Mammalian Olfactory Bulb. Anat Sci Int. 2008;83(4):218–227.

11. Otazu GH, Leibold C. A corticothalamic circuit model for sound identification in complex scenes. PLoS One. 2011;6(9):e24270. doi:10.1371/journal.pone.0024270.

12. Otazu G. Robust method for finding sparse solutions to linear inverse problems using an L2 regularization. arXiv:170100573. 2017;.

13. Boyd AM, Sturgill JF, Poo C, Isaacson JS. Cortical feedback control of olfactory bulb circuits. Neuron. 2012;76(6):1161–1174. doi:10.1016/j.neuron.2012.10.020.

14. Lepousez G, Nissant A, Bryant AK, Gheusi G, Greer CA, Lledo PM. Olfactory learning promotes input-specific synaptic plasticity in adult-born neurons. Proc Natl Acad Sci USA. 2014;111:13984. doi:10.1073/pnas.1404991111.

15. Otazu GH, Chae H, Davis MB, Albeanu DF. Cortical Feedback Decorrelates Olfactory Bulb Output in Awake Mice. Neuron. 2015;86(6):1461–1477. doi:10.1016/j.neuron.2015.05.023.

16. Livneh Y, Mizrahi A. Experience-dependent plasticity of mature adult-born neurons. Nat Neurosci. 2012;15(1):26–28.

17. Sailor KA, Valley MT, Wiechert MT, Riecke H, Sun GJ, Adams W, et al. Persistent Structural Plasticity Optimizes Sensory Information Processing in the Olfactory Bulb. Neuron. 2016;91(2):384–396. doi:10.1016/j.neuron.2016.06.004.

18. Lepousez G, Valley MT, Lledo PM. The impact of adult neurogenesis on olfactory bulb circuits and computations. Annual review of physiology. 2013;75:339–363. doi:10.1146/annurev-physiol-030212-183731.

19. Yamaguchi M, Mori K. Critical period for sensory experience-dependent survival of newly generated granule cells in the adult mouse olfactory bulb. Proc Natl Acad Sci USA. 2005;102(27):9697–9702.

20. Wallace JL, Wienisch M, Murthy VN. Development and Refinement of Functional Properties of Adult-Born Neurons. Neuron. 2017;doi:10.1016/j.neuron.2017.09.039.

21. Huang LW, Ung K, Garcia I, Quast KB, Cordiner K, Saggau P, et al. Task Learning Promotes Plasticity of Interneuron Connectivity Maps in the Olfactory Bulb. J Neurosci. 2016;36(34):8856–8871. doi:10.1523/JNEUROSCI.0794-16.2016.

22. Sultan S, Mandairon N, Kermen F, Garcia S, Sacquet J, Didier A. Learning-Dependent Neurogenesis in the Olfactory Bulb Determines Long-Term Olfactory Memory. Faseb J. 2010;24(7):2355–2363.

23. Alonso M, Lepousez G, Wagner S, Bardy C, Gabellec MM, Torquet N, et al. Activation of adult-born neurons facilitates learning and memory. Nat Neurosci. 2012;15:897.

24. Moreno MM, Linster C, Escanilla O, Sacquet J, Didier A, Mandairon N. Olfactory perceptual learning requires adult neurogenesis. Proc Natl Acad Sci U S A. 2009;106(42):17980.

25. Sultan S, Rey N, Sacquet J, Mandairon N, Didier A. Newborn Neurons in the Olfactory Bulb Selected for Long-Term Survival through Olfactory Learning Are Prematurely Suppressed When the Olfactory Memory Is Erased. J Neurosci. 2011;31(42):14893–14898.

26. Mandairon N, Kermen F, Charpentier C, Sacquet J, Linster C, Didier A. Context-driven activation of odor representations in the absence of olfactory stimuli in the olfactory bulb and piriform cortex. Front Behav Neurosci. 2014;8:138. doi:10.3389/fnbeh.2014.00138.

27. Thompson AD, Picard N, Min L, Fagiolini M, Chen C. Cortical Feedback Regulates Feedforward Retinogeniculate Refinement. Neuron. 2016;91:1021–1033. doi:10.1016/j.neuron.2016.07.040.

28. Fukunaga I, Berning M, Kollo M, Schmaltz A, Schaefer AT. Two distinct channels of olfactory bulb output. Neuron. 2012;75(2):320–329. doi:10.1016/j.neuron.2012.05.017.

29. Manabe H, Mori K. Sniff rhythm-paced fast and slow gamma-oscillations in the olfactory bulb: relation to tufted and mitral cells and behavioral states. Journal of Neurophysiology. 2013;110(7):1593–1599. doi:10.1152/jn.00379.2013.

30. Hasselmo ME. Acetylcholine and Learning In A Cortical Associative Memory. Neural Computation. 1993;5(1):32–44. doi:10.1162/neco.1993.5.1.32.

31. Barnes DC, Hofacer RD, Zaman AR, Rennaker RL, Wilson DA. Olfactory perceptual stability and discrimination. Nat Neurosci. 2008;11(12):1378–1380.

32. Chapuis J, Wilson DA. Bidirectional plasticity of cortical pattern recognition and behavioral sensory acuity. Nat Neurosci. 2011;15(1):155.

33. Mandairon N, Jourdan F, Didier A. Deprivation of Sensory Inputs To the Olfactory Bulb Upregulates Cell Death and Proliferation in the Subventricular Zone of Adult Mice. Neuroscience. 2003;119(2):507–516.

34. Lin CW, Sim S, Ainsworth A, Okada M, Kelsch W, Lois C. Genetically increased cell-intrinsic excitability enhances neuronal integration into adult brain circuits. Neuron. 2010;65:32–9.

35. Chow SF, Wick SD, Riecke H. Neurogenesis Drives Stimulus Decorrelation in a Model of the Olfactory Bulb. PLoS Comp Biol. 2012;8:e1002398.

36. Shepherd GM. Olfactory Bulb. In: Shepherd GM, editor. The Synaptic Organization of the Brain. Oxford University Press; 2004. p. 165.

37. Rokni D, Hemmelder V, Kapoor V, Murthy VN. An olfactory cocktail party: figure-ground segregation of odorants in rodents. Nat Neurosci. 2014;17(9):1225–1232. doi:10.1038/nn.3775.

38. Gschwend O, Abraham NM, Lagier S, Begnaud F, Rodriguez I, Carleton A. Neuronal pattern separation in the olfactory bulb improves odor discrimination learning. Nature Neuroscience. 2015;18:1474–1482. doi:10.1038/nn.4089.

39. Kelsch W, Lin CW, Lois C. Sequential development of synapses in dendritic domains during adult neurogenesis. Proc Natl Acad Sci USA. 2008;105(43):16803–16808.

40. Linster C, Sachse S, Galizia CG. Computational Modeling Suggests That Response Properties Rather Than Spatial Position Determine Connectivity Between Olfactory Glomeruli. J Neurophysiol. 2005;93(6):3410–3417.

41. Wick S, Wiechert M, Friedrich R, Riecke H. Pattern orthogonalization via channel decorrelation by adaptive networks. J Comput Neurosci. 2010;28:29–45.

42. Redondo RL, Kim J, Arons AL, Ramirez S, Liu X, Tonegawa S. Bidirectional switch of the valence associated with a hippocampal contextual memory engram. Nature. 2014;513(7518):426–430. doi:10.1038/nature13725.

43. Mandairon N, Sultan S, Nouvian M, Sacquet J, Didier A. Involvement of newborn neurons in olfactory associative learning? The operant or non-operant component of the task makes all the difference. The Journal of Neuroscience. 2011;31(35):12455–12460.

44. Ming Gl, Song H. Adult Neurogenesis in the Mammalian Brain: Significant Answers and Significant Questions. Neuron. 2011;70(4):687–702. doi:10.1016/j.neuron.2011.05.001.

45. Aimone JB, Li Y, Lee SW, Clemenson GD, Deng W, Gage FH. Regulation and function of adult neurogenesis: from genes to cognition. Physiological Reviews. 2014;94:991–1026. doi:10.1152/physrev.00004.2014.

46. Aimone JB, Gage FH. Modeling new neuron function: a history of using computational neuroscience to study adult neurogenesis. Eur J Neurosci. 2011;33(6):1160–1169.

47. Cecchi GA, Petreanu LT, Alvarez-Buylla A, Magnasco MO. Unsupervised Learning and Adaptation in a Model of Adult Neurogenesis. J Comput Neurosci. 2001;11(2):175–182.

48. Soucy ER, Albeanu DF, Fantana AL, Murthy VN, Meister M. Precision and diversity in an odor map on the olfactory bulb. Nat Neurosci. 2009;12(2):210–220.

49. Stettler DD, Axel R. Representations of odor in the piriform cortex. Neuron. 2009;63(6):854–864.

50. Fiser A, Mahringer D, Oyibo HK, Petersen AV, Leinweber M, Keller GB. Experience-dependent spatial expectations in mouse visual cortex. Nature Neuroscience. 2016;19:1658–1664. doi:10.1038/nn.4385.

51. Dolmetsch RE, Pajvani U, Fife K, Spotts JM, Greenberg ME. Signaling to the nucleus by an L-type calcium channel-calmodulin complex through the MAP kinase pathway. Science. 2001;294(5541):333–339.

52. Wang W, Lu S, Li T, Pan YW, Zou J, Abel GM, et al. Inducible activation of ERK5 MAP kinase enhances adult neurogenesis in the olfactory bulb and improves olfactory function. J Neurosci. 2015;35(20):7833–7849. doi:10.1523/JNEUROSCI.3745-14.2015.

53. Kay LM, Beshel J, Brea J, Martin C, Rojas-Libano D, Kopell N. Olfactory oscillations: the what, how and what for. Trends Neurosci. 2009; p. 2510–2510.

54. Linster C, Cleland TA. Decorrelation of odor representations via spike timing-dependent plasticity. Front Comp Neurosci. 2010;4:157.

55. Wilson CD, Serrano GO, Koulakov AA, Rinberg D. Concentration invariant odor coding. Nature Comm. 2017;8:1477. doi:http://dx.doi.org/10.1101/125039.

56. Isaacson JS. Mechanisms Governing Dendritic Gamma-Aminobutyric Acid (GABA) Release in the Rat Olfactory Bulb. Proc N Acad USA. 2001;98(1):337–342.

57. Egger V, Svoboda K, Mainen ZF. Dendrodendritic Synaptic Signals in Olfactory Bulb Granule Cells: Local Spine Boost and Global Low-Threshold Spike. J Neurosci. 2005;25(14):3521–3530.

58. Egger V. Synaptic sodium spikes trigger long-lasting depolarizations and slow calcium entry in rat olfactory bulb granule cells. Eur J Neurosci. 2008;27(8):2066–2075.

59. Osinski BL, Kay LM. Granule cell excitability regulates gamma and beta oscillations in a model of the olfactory bulb dendrodendritic microcircuit. J Neurophys. 2016;116(2):522–539. doi:10.1152/jn.00988.2015.

60. Nissant A, Bardy C, Katagiri H, Murray K, Lledo PM. Adult neurogenesis promotes synaptic plasticity in the olfactory bulb. Nat Neurosci. 2009;12(6):728–730.

61. Gao Y, Strowbridge BW. Long-term plasticity of excitatory inputs to granule cells in the rat olfactory bulb. Nat Neurosci. 2009;12(6):731–733.

62. Ma TF, Zhao XL, Cai L, Zhang N, Ren SQ, Ji F, et al. Regulation of Spike Timing-Dependent Plasticity of Olfactory Inputs in Mitral Cells in the Rat Olfactory Bulb. Plos One. 2012;7(4):e35001. doi:10.1371/journal.pone.0035001.

